# Pathological relevance of post-translationally modified alpha-synuclein (pSer87, pSer129, nTyr39) in idiopathic Parkinson’s disease and Multiple System Atrophy

**DOI:** 10.1101/2022.01.11.475823

**Authors:** Berkiye Sonustun, Firat M Altay, Catherine Strand, Geshanthi Hondhamuni, Thomas T Warner, Hilal A Lashuel, Rina Bandopadhyay

**Affiliations:** Reta Lila Weston Institute and Department of Movement Neuroscience, UCL Queen Square Institute of Neurology, 1Wakefield Street, WC1N 1PJ; Department of Neuroscience, Graduate School of Medical Sciences, Weill Cornell Medical College, Cornell University, NY USA; Laboratory of Molecular and Chemical Biology of Neurodegeneration, School of Life Sciences, Brain Mind Institute, Ecole Polytechnique Fédérale de Lausanne (EPFL), CH-1015 Lausanne, Switzerland; Queen Square Brain Bank, UCL Queen Square Institute of Neurology, 1Wakefield Street, WC1N 1PJ

**Keywords:** alpha-synuclein, post-translational modifications, Parkinson’s disease, Multiple system atrophy, Lewy bodies, Lewy neurites, Glial cytoplasmic inclusions, phosphorylation, nitration, immunohistochemistry

## Abstract

Aggregated alpha-synuclein (α-synuclein) is the main component of Lewy bodies (LBs), Lewy neurites (LNs), and glial cytoplasmic inclusions (GCIs), which are pathological hallmarks of idiopathic Parkinson’s disease (IPD) and multiple system atrophy (MSA), respectively. Initiating factors that culminate in forming LBs/LNs/GCIs remain elusive. Several species of α-synuclein exist, including phosphorylated and nitrated forms. It is unclear which α-synuclein post-translational modifications (PTMs) appear within aggregates throughout disease pathology. Herein we aimed to establish the predominant α–synuclein PTMs in post-mortem IPD and MSA pathology using immunohistochemistry. We examined the patterns of three α-synuclein PTMs (pS87, pS129, nY39) simultaneously in pathology-affected regions of 15 PD, 5 MSA, 6 neurologically normal controls. All antibodies recognized LBs, LNs, and GCIs, albeit to a variable extent. pS129 α-synuclein antibody was particularly immunopositive for LNs and synaptic dot-like structures followed by nY39 α-synuclein antibody. GCIs, neuronal inclusions, and small threads were positive for nY39 α-synuclein in MSA. Quantification of the LB scores revealed that pS129 α-synuclein was the dominant and earliest α-synuclein PTM followed by nY39 α-synuclein, while lower amounts of pSer87 α-synuclein appeared later in disease progression in PD. These results may have implications for novel biomarker and therapeutic developments.

## 1. Introduction

Parkinson’s disease (PD) is the second most prevalent chronic progressive neurodegenerative disorder, affecting more than 1% of population above 60 years of age [1]. Along with typical motor dysfunctions of bradykinesia, rigidity, and rest tremor, patients often manifest a range of non-motor symptoms from anosmia to rapid eye movement (REM) sleep disorder, constipation and depression [2, 3]. Whilst familial forms of the disease have been identified [4] with mutations in the alpha-synuclein (α-synuclein) gene found as the first causal link associating families with autosomal dominant PD, the majority (around 90%) of the cases remain of unknown origin. Although still not well understood, various genetic risk factors have also come to light in the last two decades through genetic and Genome-Wide Association Studies [5, 6] that may associate with idiopathic PD (IPD), including *SNCA* and *LRRK2*.

Degeneration of the dopaminergic nigrostriatal system is a prominent pathological feature of PD, leading to impaired dopaminergic neurotransmission within the basal ganglia. The presence of aggregated α-synuclein within cytoplasmic Lewy bodies (LBs) and dystrophic Lewy neurites (LNs) are also common pathological features at post-mortem [7]. Braak et al. [8] have proposed a model wherein it is suggested that LB pathology in PD arises in the dorsal motor nucleus of the vagus or the anterior olfactory nucleus before affecting the nigra and the limbic regions, followed by spreading in higher cortical regions [8]. Increasing evidence suggests that this occurs via the putative prion-like spread of α-synuclein [9].

Multiple system atrophy (MSA) is a rare but rapidly progressing neurodegenerative disorder of uncertain etiology. Currently, there are no disease-modifying therapies available for MSA. It is clinically characterised by parkinsonism, cerebellar, and motor dysfunctions [10]. The neuropathological hallmark lesion of MSA features mainly in the oligodendroglia [glial cytoplasmic inclusions (GCIs)] and are immunoreactive for α-synuclein. Less frequently, cytoplasmic inclusions (NCIs) and neuronal nuclear inclusions (NNIs) are observed in some anatomical regions together with neuronal threads, which are all α-synuclein immunopositive. Although the brunt of the neuronal loss is observed in the striatonigral and olivopontocerebellar regions, the neurons of locus coeruleus and the dorsal vagal nuclei are also affected [10,11].

Common post-translational modifications (PTMs) such as phosphorylation and nitration of proteins can occur in disease pathogenesis, and α-synuclein is known to undergo varied and extensive PTMs [reviewed in 12,13]. These covalent PTMs may play a role in protein folding and intraneuronal aggregation and propagation through mechanisms that modify its conformational landscape, membrane association, degradation, and/or interactome [14]. The C-terminus of α-synuclein plays a critical role in regulating the interactions of α-synuclein with other proteins and ligands such as calcium, polyamines, dopamine, and metal ions [15]. Notably, the majority of disease-associated PTM sites in α-synuclein are located in the C-terminus, implying that these modifications may be involved not only in the regulation of structure and physiological function of α-synuclein, but also its aggregation, pathology formation, and spreading.

Previous studies have demonstrated that α-synuclein is constitutively phosphorylated at different residues [16] and that phosphorylation at Ser129 (pS129) residue is the dominant PTM of α-synuclein within LBs, LNs, and GCIs [17]. Furthermore, biochemical fractionation from post-mortem brains also reported that over 90% of the insoluble α-synuclein found in dementia with Lewy body (DLB) cases are phosphorylated at S129 compared to the 4% seen in healthy brains [17], thereby implicating phosphorylation at this residue as a potential key event in α-synuclein pathology formation, spreading or clearance. Additionally, both α-synuclein within inclusions in glia and neurons of MSA brains [18] are immunoreactive to pS129 α-synuclein antibodies. Although phosphorylation at S129 is robustly associated with α-synuclein inclusion formation in several synucleinopathies, the mechanisms by which this or other PTMs influence α-synuclein aggregation and contribute to Lewy pathology formation and spreading in the brain remain unclear.

In contrast to pS129 α-synuclein, pS87 α-synuclein’s role in disease remains to be elucidated. This serine residue is found in a hydrophobic stretch of the protein’s non-amyloid component (NAC) region, which may be essential for aggregation [19]. Additionally, the presence of a charged phosphate group can potentially impact the protein structure, its oligomerisation, and its function [19]. Furthermore, Paleologou et al 2010 [20] also reported higher expression of pS87 in Alzheimer’s disease (AD), MSA, and DLB, relative to healthy controls. Previously, one study proposed a neuroprotective role of this PTM [21]. Interestingly, phosphorylation of S87 residue has been shown to exert strong aggregation inhibitory effects via increasing the conformational flexibility of α-synuclein and decreasing its affinity for lipid membranes and vesicles [20].

Markers of oxidized proteins, lipids, and DNA are upregulated in dopaminergic (DA) neurons of PD patients [22], suggesting increased levels of oxidative stress. Substantia nigra DA neurons are particularly susceptible to oxidative injury and appear to have a greater output of reactive oxygen species (ROS) [23]. Previously, Duda et al [24] demonstrated an abundance of nitrated forms of α-synuclein in LBs, LNs, and GCIs in human post-mortem brains. Further studies have indicated Y39, Y125, Y133, and Y136 to be the tyrosine nitration sites within α-synuclein. nY39 α-synuclein is found to form morphologically distinct fibrils relative to WT α-synuclein and show less affinity to negatively charged vesicles [25]. This is important, as the physiological function of α-synuclein is thought to arise through its interaction with lipid bilayers at the presynaptic terminal, to regulate synaptic vesicle docking and fusion [26]. Although WT α-synuclein aggregated faster than nY39 α-synuclein *in-vitro*, the latter was shown to form shorter and wider aggregates [25], pointing to a role of PTMs in regulating fibril strains and/or the formation of Lewy pathologies. The introduction of a negatively charged group at Y39 (i.e., phosphorylation or nitration) was previously shown to decrease the binding capacity of a nitrated α-synuclein mutant to negatively charged vesicles [25, 27]. For this study, our focus was nitration of Y39.

Herein, we have investigated the differential distribution and abundance of some key post-translational modifications of α-synuclein in IPD and MSA post-mortem brains. We aimed to correlate the appearance of PTMs with the development of the disease by assessing the α-synuclein PTMs in brain regions that are affected at different stages of the disease. Specifically, we used immunohistochemistry to study the comparisons between pS87, pS129, and nY39 α-synuclein and unmodified α-synuclein [(α-synuclein (UN)] in IPD and MSA. The differential expressions of pS87, pS129, and nY39 α-synuclein have not previously been studied in tandem in human post-mortem brains of IPD and MSA. Moreover, pS87 α-synuclein and nY39 α-synuclein have been less extensively studied relative to pS129 α-synuclein. We report that pS129 α-synuclein is a major modification in IPD and MSA, followed by nY39 α-synuclein, and finally the least numbers of LBs/GCIs positive for pS87 α-synuclein were noted in the two diseases. To our knowledge, this is the first report on the expression of nY39 α-synuclein using a novel antibody in human post-mortem IPD and MSA brains.

## 2. Material and Methods

### 2.1 Source of Brain Tissue

Human brain tissue was obtained from the Queen Square Brain Bank for Neurological Disorders archives, in which collection was done with ethical approval from the London Multicentre Research Ethics Committee UCL IoN HTA License #12198. Brain tissue from 15 IPD, 5 MSA, and 6 neurologically normal controls were examined. Limited patient demographic data are presented in Table 1.

**Table 1:** Selected demographics of the cases used.

| Case (Patient (P)/control(C)) | Gender (M/F) | Age at death (years) | Post-mortem delay (hrs:min) | Diagnosis |
| --- | --- | --- | --- | --- |
| P | F | 83 | 19:00 | IPD BStem |
| P | M | 81 | 51:00 | IPD BStem |
| P | F | 83 | 99:00 | IPD BStem |
| P | M | 74 | 19:05 | IPD BStem |
| P | M | 75 | 29:31 | IPD BStem |
| P | M | 84 | 75:05 | IPD Limbic |
| P | F | 80 | 39:10 | IPD Limbic |
| P | M | 78 | 54:55 | IPD Limbic |
| P | M | 93 | 82:20 | IPD Limbic |
| P | F | 72 | 79:40 | IPD Limbic |
| P | M | 72 | 100:20 | IPD Neocortical |
| P | M | 81 | 33:25 | IPD Neocortical |
| P | M | 81 | 73:15 | IPD Neocortical |
| P | F | 76 | 87:05 | IPD Neocortical |
| P | F | 76 | 76:15 | IPD Neocortical |
| P | F | 80 | 23:20 | MSA |
| P | M | 50 | 6:15 | MSA |
| P | F | 65 | 10:25 | MSA |
| P | F | 60 | 9:30 | MSA |
| P | M | 69 | 43:15 | MSA |
| C | M | 76 | 76:00 | Control |
| C | F | 84 | 84:00 | Control |
| C | M | 80 | 80:00 | Control |
| C | M | 83 | 83:15 | Control |
| C | F | 91 | 91:00 | Control |
| C | M | 76 | 76:00 | Control |

### 2.2. Immunohistochemistry

Paraffin embedded sections of 8μm thickness were cut from different brain regions using a microtome. Sections were de-waxed in xylene followed by treatment with 0.3% H2O2 in 100% methanol for 10 minutes to block endogenous peroxidase reactions. Sections were pretreated with 98% formic acid at room temperature for 10 minutes, followed by pressure cooking in Citrate Buffer (pH:7.0) for 10 minutes. Following a series of washes using 1 x TBS Tween-20 buffer, sections were blocked with 10% Dried Skimmed Milk Powder (Marvel) in 1 x TBS Tween 20 for 30 minutes. 4 different α-synuclein [α-synuclein UN (C-terminal); phosphorylated α-synuclein Ser87 (pS87), phosphorylated α-synuclein Ser129 (pS129), and nitrated α-synuclein Tyrosine 39 (nY39)] primary antibodies were used at 1:500 dilution for the incubation times between (1hr to overnight) and is detailed in Table 2. After a series of thorough washes with 1 x TBS Tween-20, the slides were probed with biotinylated secondary antibody for 30 minutes at room temperature before being treated with Avidin-Biotin-Complex for another 30 minutes at room temperature. After this, the slides were treated with hydrogen peroxide (0.03%), activated 3’3’-Diaminobenzidine, and counterstained by Meyer’s haematoxylin dye. Subsequently, the slides were dehydrated in graded ethanol concentrations (70-100%), cleared in xylene before permanent mounting in DPX (BDH), and coverslips for microscopy. The pS87 staining was performed using the automated stainer (Menarini) protocol.

**Table 2:** List of antibodies used in the study

| Antibodies | Source /Cat no or clone | Host and Clonality | Antibody dilution used | incubation time |
| --- | --- | --- | --- | --- |
| anti-alpha-synuclein (epitope: aa100 to C-terminal) | Abcam /ab15530 | Rabbit Polyclonal | 1 in 500 | o/n at 4C |
| anti-alpha-synuclein (phosphorylated S87) | Hilal Lashuel/Lash-pS87 | Rabbit polyclonal | 1 in 500 | 1 hr at RT |
| anti-alpha-synuclein (phosphorylated S129) | Biolegend / P-syn-81A | Mouse monoclonal | 1 in 500 | 1hr at RT |
| anti-alpha-synuclein (nitrated Y39) | Hilal Lashuel/Lash-EGT-nY39 | Rabbit Polyclonal | 1 in 500 | o/n at 4C |

### 2.3 Immunohistochemistry analysis and pathology grading in PD and MSA

Immunohistochemistry was performed on various anatomical regions of PD and MSA brains. Specifically, for PD, we examined the medulla and pons (early affected regions), the substantia nigra and cingulate cortex (regions affected mid-way during pathology spread), frontal, parietal, and temporal cortices (late affected regions), and for MSA, the medulla, pons, substantia nigra, and the cerebellum were assessed for pathology. These were then analysed using a light microscope at a low power-field (x20). To keep the observations uniform and standardized, the neuroanatomical region of interest (ROI) for each region was demarked on the slides, which were kept uniform throughout the whole analysis process. The criteria for LB and LN pathology of Lewy Body pathology based on McKeith Criteria (Adapted from McKeith et al., 2005 [28]) observed in a low power field (x20) versus the corresponding grade and severity of pathology is summarised in Tables 3. Also, we have analysed the different types of MSA pathology, i.e., GCIs, NCIs, threads were noted in the different neuroanatomical regions with the PTM α-synuclein antibodies.

**Table 3:** Lewy Body grading used in the study

| Number of Lewy bodies in X20 field | Corresponding grade of pathology |
| --- | --- |
| 0 | None |
| 1 | Grade 1 (Mild) |
| 2-5 | Grade 2 (Moderate) |
| 6-9 | Grade 3 (Severe) |
| >10 | Grade 4 (Very severe) |

### 2.4 Statistical analysis methods and representation of data

Statistical analyses on differences in outcome measures were undertaken using a non-parametric Kruskal-Wallis ANOVA test with Dunn’s multiple comparisons as a post-hoc to compare the differences between α-synuclein PTMs (versus controls) for immunohistochemical studies. A Chi-Square test of independence was also used for immunohistochemistry data to check the relative pathology frequencies in different PD groups (and controls) using our LB-Pathology grading protocol as detailed in Table 3. Mean ± SEM was noted as a measure of dispersion, and a p-value of <0.05 was considered statistically significant.

## 3. Results

### 3.1 a-synuclein pathology in PD

We sought to characterize the staining patterns of our PTM antibodies (pS87, pS129, nY39 relative to unmodified (UN) α-synuclein in human PD, MSA and neurologically healthy control brains. The epitope of α-synuclein (UN) antibody was specific for the C-terminal according to the information on the company website (refer to Table2). We observed positive immunoreactivity with all the α-synuclein antibodies examined in all IPD and MSA cases. We did not observe any immunoreactivity with α-synuclein (UN) or the three PTM α-synuclein antibodies examined here in neurologically normal control brain tissue (Fig S1). Also, there was no appreciable staining when the primary antibodies were omitted from representative sections (Fig S2).

### 3.2 Unmodified α-synuclein (UN)

The α-synuclein (UN) antibody detected both LBs and LNs as pathological aggregates in all the PD cases (Fig 1 and 2). In PD brains, α-synuclein (UN) expression was observed in small dot-like structures within neuronal perikarya, which could represent synaptic α-synuclein. Neocortical-diffuse PD cases showed most α-synuclein (UN) pathology when normalized to the neurologically healthy control group. Brainstem-predominant PD patients showed the least α-synuclein (UN) expression in all regions. However, “fine Lewy neurites” (FLNs) were more predominant in the brainstem-predominant PD cases with α-synuclein (UN) antibody (Table 4). Both FLN and “thick Lewy neurites” (TLNs) were seen dominantly in nigra and cingulate regions of the limbic and neocortical PD cases with the α-synuclein (UN) antibody.

**Fig 1:**
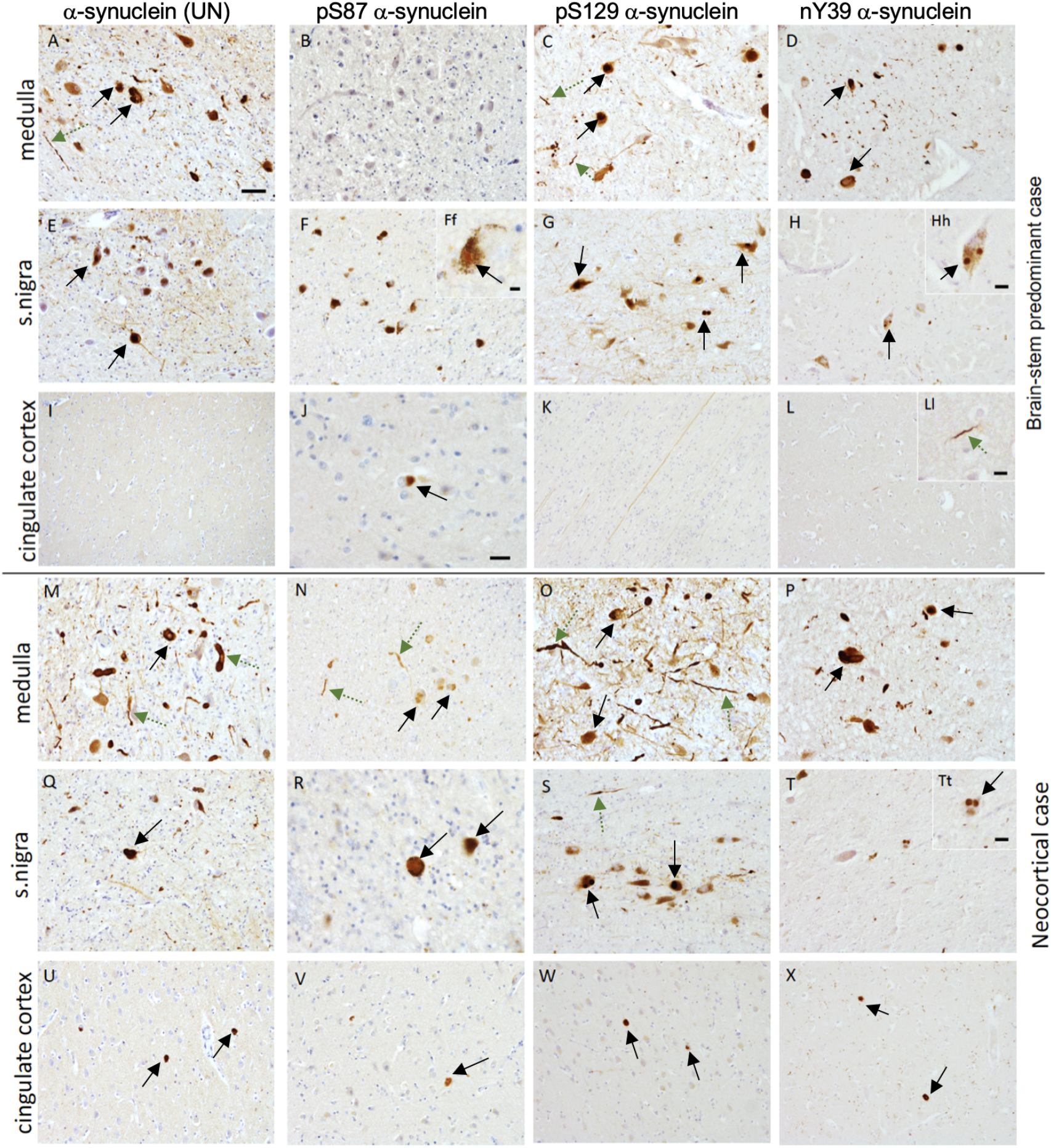
Illustration of immunostaining for different α-synuclein antibodies in IPD cases. A-L depicts regions from a brain-stem predominant PD case; A,E,I illustrates immunostaining with α-synuclein(UN) showing LBs (black arrows) and LNs (green dashed arrows) in medulla, nigra but not in cingulate cortex. Very few LBs or LNs were positive for pSer87α-synuclein, as shown in B,F,J. Positive immunostaining of LBs and LNs in medulla and s nigra but not in cingulate observed with pSer129 α-synuclein antibody. Some non-specific staining of white matter tracts were observed (K); Immunopositivity in LBs and LNs seen for nY39 α-synuclein antibody in the medulla (D) and nigra (H and Hh) and some LNs (L and Ll) in the cingulate cortex. M-X depicts regions from a neocortical PD case where higher numbers of LBs and LNs were immunopositive with all 4 antibodies. Scale Bar: 20μm in all except in J where it is 10μm. Scale bar in insets Hh, Ll, and Tt is 8μm.

**Fig 2:**
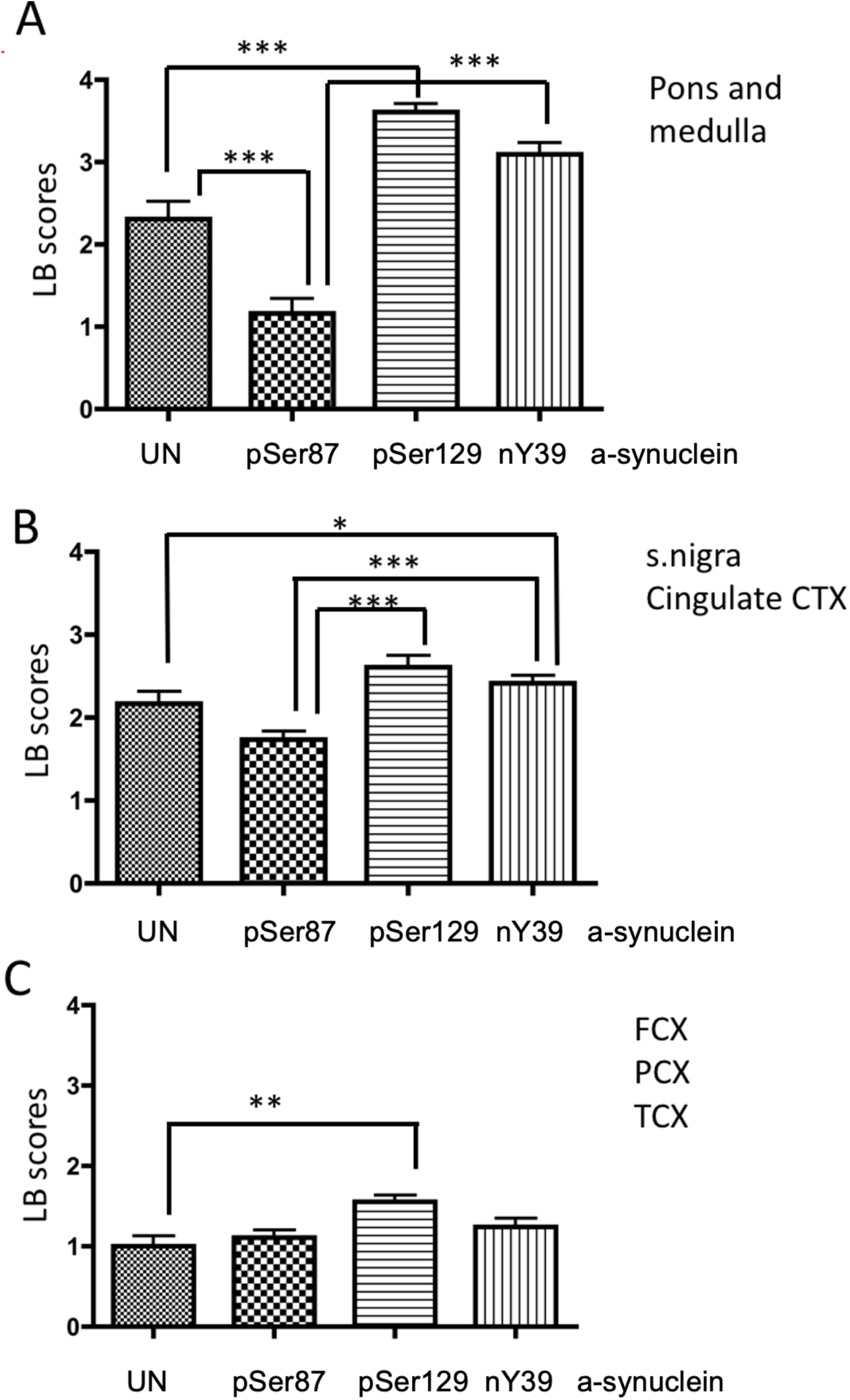
Histogram of cumulative LB scores of different α-synuclein antibodies seen in pons and medulla A); substantia nigra and cingulate cortex (CTX) B) and frontal, parietal and temporal cortex C). Statistical analysis was performed using non-parametric Kruskal-Wallis test with Dunn’s multiple comparison corrections. ***,**,* denotes p<0.001, p<0.01, p< 0.05 respectively.

**Table 4:**
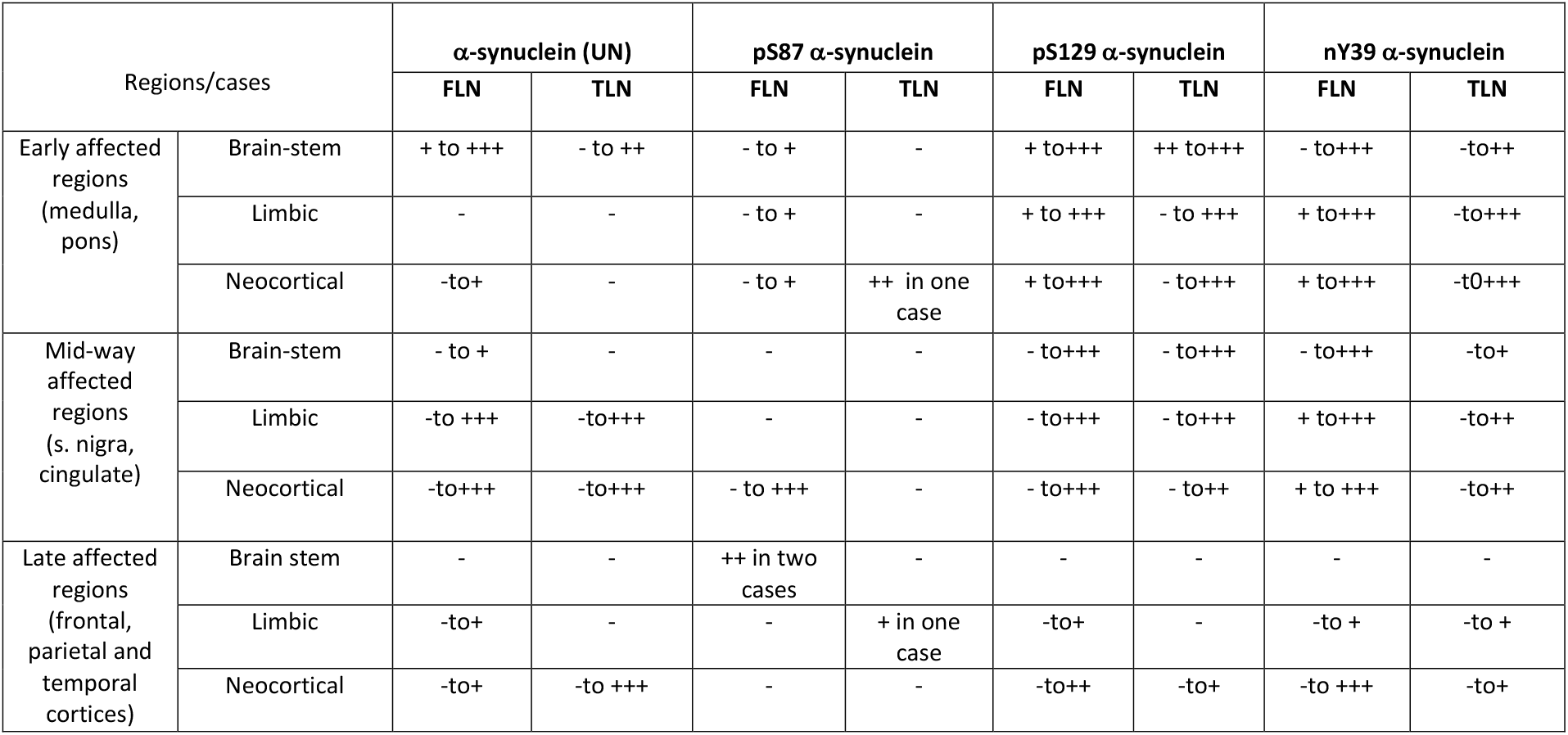
Semiquantitative assessment of Lewy neurites in IPD cases. Key to scores: - = none; +=mild; ++= moderate; +++= severe.

It is important to note that the unmodified α-synuclein (UN) antibody (epitope described as aa100 to C-terminal; Table 2) we used in our study is reported to highlight the unmodified versions of α-synuclein. The premise of our study was to compare the abundance and localization of post-translationally modified α-synuclein relative to the unmodified versions in post-mortem tissue. However, we cannot rule out whether this antibody detected C-terminal truncated species, which are known to exist in human post-mortem disease pathology. Thus, we have used α-synuclein (UN) antibody as a control to compare the modified α-synuclein antibody versions to. This was apparent from the fact that our α-synuclein (UN) antibody did not detect all of the LBs or LNs, but indeed detected dot-like structures in the neuropil, which we defined as synaptic, physiological α-synuclein.

### 3.3. pS87 a-synuclein

We demonstrate the fewest number of aggregates with pS87 α-synuclein antibody in all regions studied examined. The pS87 α-synuclein mainly recognized LB inclusions over LNs. Overall, we observed less dense staining with paler inclusions with pS87 α-synuclein antibody (Fig 1). We observed fewer amounts of pSer8 α-synuclein FLNs and TLNs in all of the IPD cases examined (Table 4).

### 3.4 pS129 α-synuclein

The pS129 α-synuclein antibody showed immunoreactivity in the early affected regions in PD cases, including many LBs and pale body-like inclusions (Fig 1 and 2). We observed dot-like structures prominently in the midbrain with the pS129 α-synuclein antibody. The nigra showed the highest pS129 α-synuclein pathology compared to other regions regardless of the disease severity (Fig 1). Moreover, this antibody recognized subcortical fibres and axons in IPD cases and also in neurological-control cases which most likely reflects non-specific neurofilament staining. Most pS129 α-synuclein pathology was observed in the nigra of the brainstem-predominant IPD group, thereby reinforcing the notion that this modification is an early event in IPD pathogenesis. The pS129 α-synuclein antibody also recognized both FLNs and TLNs, specifically in the early and mid-way affected regions (Table 4).

### 3.5 nY39 α-synuclein

This novel antibody recognized both classic nigral and cortical LBs in the subcortical and cortical regions, respectively (Fig1, 2). We observed diffuse synaptic nY39 α-synuclein immunoreactivity in the grey matter of all PD cases. High staining of FLNs and TLNs were noted for brainstem, limbic and neocortical cases in the pons, medulla, substantia nigra and cingulate regions (Table 4). Fewer neuritic staining pattern was observed in cortical regions (frontal, parietal and occipital regions) with nY39 α-synuclein compared to subcortical regions. It is noteworthy that this antibody preferentially marked FLNs in the early affected and midway affected regions (Table 4).

### 3.6 Differential co-occurrence of the α-synuclein PTMs in Lewy pathologies

#### 3.6.1 LBs

Cumulative LB scores are shown for early, mid-way and late affected anatomical brain regions in Figure 2. The differences between α-synuclein PTMs were tested using the Kruskal-Wallis test with multiple-comparisons. The LB grading scheme used for the purpose of this study is summarized in Table 3.

Our data suggest differential distribution of α-synuclein PTMs in LBs of early, mid-way, and late-affected PD regions. In the early-affected regions, pS129 α-synuclein was significantly higher in LBs, followed closely by nY39 α-synuclein compared to those with pS87 α-synuclein and α-synuclein (UN). Based on the immunoreactivity of the pS129 and nY39 α-synuclein antibodies, the most pathologically affected regions were the pons and medulla, with fewer LBs detected in the cortical regions examined in this study, namely, frontal, temporal and parietal cortices (Fig 2). In the brainstem, i.e., “early-affected region”, pS129 α-synuclein was the most abundant α-synuclein species (Fig 2A, p<0.01). We noted significant elevation of α-synuclein (UN) immunoreactivity over pS87 α-synuclein in the pons and medulla. The pS87 α-synuclein immunoreactivity was significantly lower than the other two α-synuclein PTMs in midway-affected regions (Figure 2B). In midway-affected PD regions, Grade 4 pathology with pS129 α-synuclein was relatively less in comparison to early affected regions (Fig 2A, B). We also demonstrate that pS129 α-synuclein significantly dominated over α-synuclein (UN) in early, mid-way and late-affected regions (Figure 2A, B, and C, respectively). The overall LB frequency observed in early affected regions varied between severe and very severe categories. Our data indicate that early affected PD regions are heavily affected by phosphorylated and nitrated forms of α-synuclein examined here. The amount of PTM positive inclusions diminished as the disease progressed from early to late stages.

#### 3.6.2 LNs

The semiquantitative scores of FLNs and TLNs are presented in Table 4. The LNs were categorized as FLNs and TLNs. High numbers of LNs were immunopositive for pS129 and nY39 α-synuclein antibodies. These appeared in higher numbers in early and mid-way affected regions. In comparison, pS87 detected the least number of LNs, and where it was present, these were noted predominantly in FLNs. The medulla, nigra, and cingulate gyrus demonstrated the most frequent LNs compared to other regions examined. The α-synuclein (UN) antibody detected lower numbers of both FLNs and TLNs compared to pSer129 and nY39 α-synuclein antibodies which might be due the antibody (epitope; discussed earlier in section2.2 and Table 2) not recognizing all of the α-synuclein species in post-mortem tissue. The peak levels of nY39 α-synuclein pathology were observed in the early affected regions, especially in the neocortical and diffuse-limbic PD cases. We also noted overall that the predominant α-synuclein PTM in the FLNs was nY39. However, it is noteworthy that pS129 α-synuclein detected more TLNs compared to nY39 α-synuclein, especially in the mid-way affected regions of the brainstem and limbic PD. Moreover, pS129 and nY39 α-synuclein were equally distributed among the FLNs in midway affected regions (Table 4).

#### 3.7 α-synuclein pathology in MSA

GCIs were the dominant pathogenic structures observed in all MSA cases/regions examined (Fig 3). Neuronal inclusions were also seen in all the MSA cases, prominently in the pons. NCIs were differentiated from glial inclusions as they appear larger with ovoid shape. NNIs appeared as a floating network of filaments in the nucleoplasm (Fig 3).

**Fig 3:**
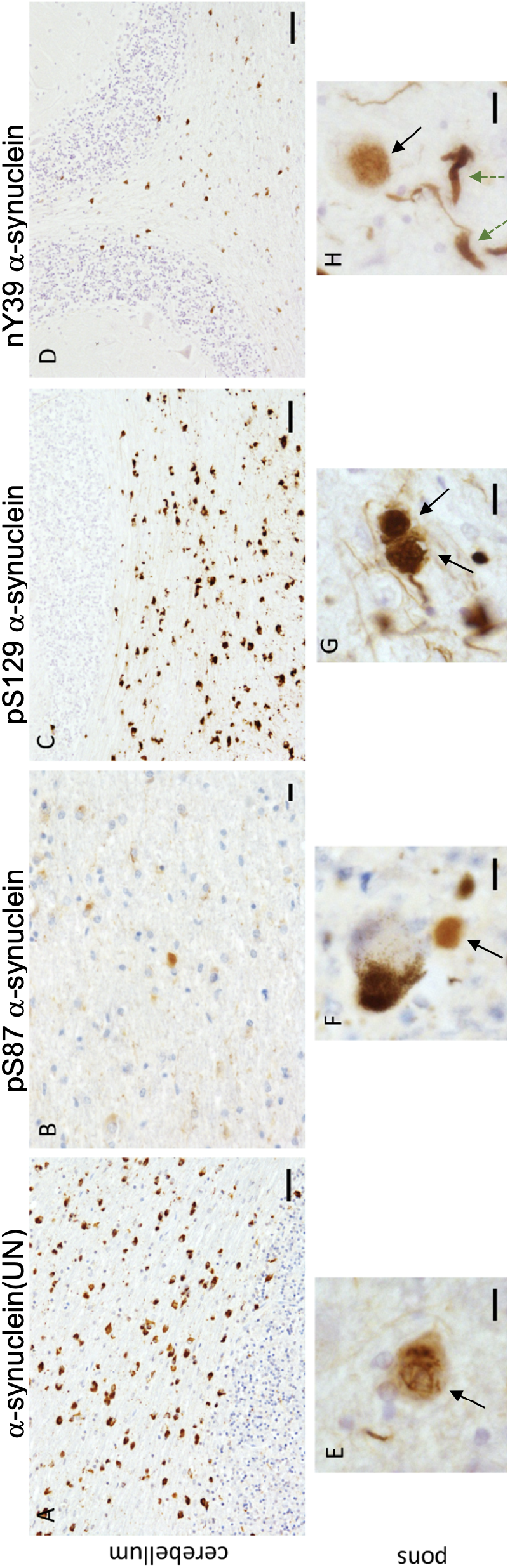
Illustration of immunostaining for different α-synuclein antibodies in cerebellum and pontine base of MSA cases. Immunostaining with α-synuclein unmodified antibody depicting numerous GCIs in the cerebellar white matter in A); and neuronal nuclear inclusion and Neuronal cytoplasmic inclusion (black arrow) in E); Few GCI’s positively stained for Ser87P α-synuclein in CBM B) and extracellular staining in pontine base F); Pser129 α-synuclein marks several GCIs in Cerebellum C) and NCIs (black arrow) and fine neurites in pontine base G); nY39 α-synuclein labels fewer GCIs in cerebellum D) and NCI’s (black arrow) and thick (green dashed arrows) and thin neurites in H). Scale bars denote 20μm in A, C, D; 10μm in B; 8μm in E, G; 5μm in F and H.

The staining patterns observed in MSA cases were akin to those observed in PD in the context of the different antibodies we have used. For instance, pS87 α-synuclein antibody demonstrated the palest-appearing aggregates in the cerebellum (Figure 3B). pS87 α-synuclein was dominantly observed in GCIs in both pons and cerebellum, accompanied by some threads in some of the cases. In MSA, neuronal inclusions were very rarely detected with this antibody, implying that this modification may not play a role early on in disease [22]. Likewise, pS129 α-synuclein staining was denser, wherein this antibody preferentially recognized both neuronal and glial inclusions, as well as threads in the cerebellum and pons (Figure 3B, 3G). Alongside, we observed more thread-like structures and some neuronal inclusions with nY39 α-synuclein antibody in the pons (Figure 3H), compared to the cerebellum where the dominant aggregates were GCIs (Figure 3D). In particular, a higher density of pathology was demonstrated in the medulla and pontine regions in the MSA cases, mainly in GCIs with α-synuclein (UN) and pS129 α-synuclein antibodies. Lower numbers of GCIs were immunopositive for nY39 α-synuclein in the all the regions examined compared to α-synuclein (UN) and pS129 α-synuclein. The pS129 α-synuclein antibody demonstrated fewer thread-like structures in MSA cases. The staining with pS129 α-synuclein was profound, and subcortical axonal fibers were recognized in the same manner as in IPD cases.

## 4. Discussion

PD and MSA demonstrate abundant aggregated deposition of the presynaptic protein α-synuclein in neurons and glial cells. The nature of the biological triggers that initiate α-synuclein aggregation has been a matter of intense research over the past 20 years. Numerous α-synuclein post-translational modifications have been identified to date, including phosphorylation, nitration, ubiquitination, acetylation, sumoylation and specific truncations [29], and some of these are key markers of disease pathogenesis. With the aim of establishing an in-depth disease-, region- and cell type-specific distribution of α-synuclein PTMs, it is important to examine these in parallel within a subset of disease cases. Therefore, in the current study, our aim was to determine the presence and relative abundance of three different α-synuclein PTMs in IPD and MSA pathology and investigate how they associate to the region and cell-type specificity at different stages of disease. For IPD, we demonstrate that the PTMs of α-synuclein display variable abundance at different anatomical sites within LBs and LNs, thereby implying different roles which interplay at different time points during pathogenesis. For MSA, we demonstrate the presence of all three PTMs (pS87, pS129, nY39) of α-synuclein examined here dominantly in GCIs but they also localized to NCIs and NNIs, suggesting that the modifications are part of disease pathology (summarised in Table 5).

**Table 5:** Summary of pathological aggregates stained with the various α-synuclein antibodies.

LBs are thought to be the basis of neurotoxicity and cell death. Nonetheless, there are opposing views in the field. The dystrophic neuronal processes, or LNs appear earlier during the pathogenesis timeline, as α-synuclein aggregates in the axons [30]. Aggregated LNs are believed to compromise neuronal function through disrupting axonal transport (reviewed in Perlson et al.,2010 [31]). It is likely that the perikaryal LBs form in an attempt to “sweep up” the surrounding LNs into a single inclusion as a counteractive mechanism, such that synaptic function can be restored. However, if this assumption is true, the point at which LBs become pathogenic, and the mechanisms underlying this process remain elusive. Curiously, we saw neither dot-like synaptic α-synuclein, nor many neurites positive for pS87 α-synuclein. In contrast, pS129 α-synuclein and nY39 α-synuclein appear to be the dominant PTMs in both LNs and LBs, and are also present in dot-like structures in the neuropil. It is likely that α-synuclein aggregates that form LNs already have these PTMs, which later amass into LBs.

In LBs, a high percentage (~90%) of α-synuclein is phosphorylated at S129 [17, 32] although whether this happens before LB formation remains a matter of debate. Recently, using a variety of sophisticated electron microscopy techniques and cryoEM it was demonstrated that LBs are not only composed non-fibrillar α-synuclein but they are also enriched with lipids and membranous components and organelles such as mitochondria and vesicles [33]. Subsequently in an *in vitro* based study, Mahul-Mellier et al [34] demonstrated that LB formation is a result of a complex interplay of α-synuclein fibrillisation, post-translational modifications together with its interactions with membranous organelles like the mitochondria and the vesicular components of the autophagosome and endolysosomes. Specifically, this study recognised pS129 α-synuclein as an early PTM that could also regulate other α-synuclein PTMs such as ubiquination and C-terminal truncation. Our data suggest that pSer129 is a dominant PTM observed in IPD and this may favour LB development.

In a previous study using the same antibody we used, Paleologou et al showed that pSer87α-synuclein levels are increased in synucleinopathies [20]. In our study we show that pS87 immunoreactivity was present in a proportion of both cortical and nigral LBs. pS87 α-synuclein was also seen in a small number of LNs in some early affected regions and sparse TLNs. This is suggestive that phosphorylaton at S87 occurs later than pS129 which is concordant with previous studies demonstrating that phosphorylation at S87 significantly inhibits the aggregation of monomeric a-synulcein [20].

Nitration is an interesting PTM, as it has a direct relationship with oxidative stress and injury. The presence of nitro-tyrosine has been demonstrated in the vicinity of oxidative injury in LB-bearing neurons [35]. Indeed, it was suggested that nitration may in fact be the initiating event in aggregate formation in LBs [36]. Peroxynitrite, both an oxidating and nitrating agent, appears to target α-synuclein, wherein nitrated α-synuclein was shown to accelerate fibrillation process [37]. We have shown that nY39 α-synuclein positivity in LBs were higher in early and mid-way affected regions with the peak being observed in pontine and medullary LBs. Lower numbers of cortical LBs were immunopositive for nY39 α-synuclein. This may suggest that nY39 is an early event or nitration of α-synuclein affects subcortical neurons more specifically. With regards to its presence in LNs, nY39 predominantly immunolabelled FLNS over TLNS, again implying that nY39 α-synuclein arises early on in IPD pathogenesis.

Alpha-synuclein positive cytoplasmic inclusions or GCIs are specific pathological hallmarks of MSA. The source of α-synuclein in oligodendrocytes remain enigmatic, although some studies have suggested neuronal α-synuclein internalisation through endocytosis, enhanced expression and decreased degradation of oligodendroglial α-synuclein as plausible theories [reviewed in 38]. We demonstrate that unmodified α-synuclein, pS87, pS129 and nY39 α-synuclein were all present in GCIs albeit to a variable extent. Additionally, all three α-synuclein modifications examined here were also associated with NCIs and NNIs suggesting that all three α-synuclein PTMs are integrated unevenly in their formation although when and why they appear in these inclusions in MSA pathology is a matter of debate [11, 18].

Our results validated that pS129 α-synuclein represents the most NCI and NNI pathology frequency compared to other PTMs, suggesting that this PTM occurs early in MSA pathogenesis. The pS87 α-synuclein antibody demonstrated the palest-appearing MSA aggregates compared to the rest of the antibodies. Moreover, pS87 modification was dominantly observed in glial inclusions, accompanied by some threads in some of the cases. Few NNIs were detected with pS87, implying that it does not represent an early-type PTM. Likewise, pS129 α-synuclein staining was more intense and detected in neuronal as well as glial inclusions and threads. Nevertheless, the dominant inclusions associated with this PTM were the GCIs, which were unambiguously present in the cerebellar white matter, pontine base and medulla but to a lesser extent in the nigral regions. The most frequent threads were observed with nY39 α-synuclein, mainly in the pons. GCIs were sparsely detected with nY39 α-synuclein compared to pS129 α-synuclein, suggesting that this modification could occur later in disease pathogenesis in MSA, most likely due to higher levels of oxidative stress. In comparison to nY39 α-synuclein, pS129 α-synuclein demonstrated fewer threadlike structures in MSA cases. This antibody also demonstrated some synaptic α-synuclein staining, which implies that pS129 is a modification that occurs dominantly in pathological scenarios and may also be upregulated in synapses in MSA consistent with the literature [39].

## 5. Conclusion

We conclude that various post-translationally modified forms of α-synuclein exist in PD and MSA-related inclusions. In both diseases, pS129 α-synuclein and nY39 α-synuclein were present in pathological aggregates, mainly presenting in early-type inclusions. pS129 α-synuclein appears to be the dominant and earliest α-synuclein PTM, while pS87 α-synuclein appears later in disease progression in IPD. Similarly, pS129 α-synuclein is the dominant PTM in MSA. We also demonstrate for the first time the presence of nY39 α-synuclein in IPD and MSA pathological inclusions. This current study extends the array of α-synuclein PTMs in the context of disease pathologies in IPD and MSA. We acknowledge that the study is limited by the use of small number of cases however, it forms a platform for a deeper understanding of α-synuclein modifications and its pathological relevance in two key α-synucleinopathies; IPD and MSA. Clearly, larger case-cohort studies are warranted including validation and quantification using other techniques such as specific ELISAs. A recent study has elegantly demonstrated diversity of α-synuclein C-terminal truncations in discriminating different synucleinopathies [40]. By continuing to interrogate human post-mortem tissue pathology using novel α-synuclein antibodies, our understanding of disease pathogenesis will increase profoundly and may also pose as a basis of both biomarker (prognostic and diagnostic) and therapeutic discoveries.

## Acknowledgements

RB is funded by the Reta Lila Weston Trust Fund and British Neuropathological Society. HAL and FMA are funded by the EPFL and the Michael J Fox foundation.

## Table and Figure Legends

Fig S1: Representative immunohistochemistry illustrations of neurologically normal control cases showing no abnormal α-synuclein deposits when treated with various α-synuclein antibodies. Scale bar 50μm in all.

Fig S2: Immunohistochemistry of brain sections from IPD cases showing no appreciable staining within PD nigra (A) or frontal cortex (B) when primary antibodies were omitted but treated with anti-rabbit and anti-mouse secondaries respectively. Scale bar 60μm in A, 15μm in B.

