## Supplementary Figs 1 and 2. for "Pathological relevance of post-translationally modified alpha-synuclein (pSer87, pSer129, nTyr39) in idiopathic Parkinson’s disease and Multiple System Atrophy"

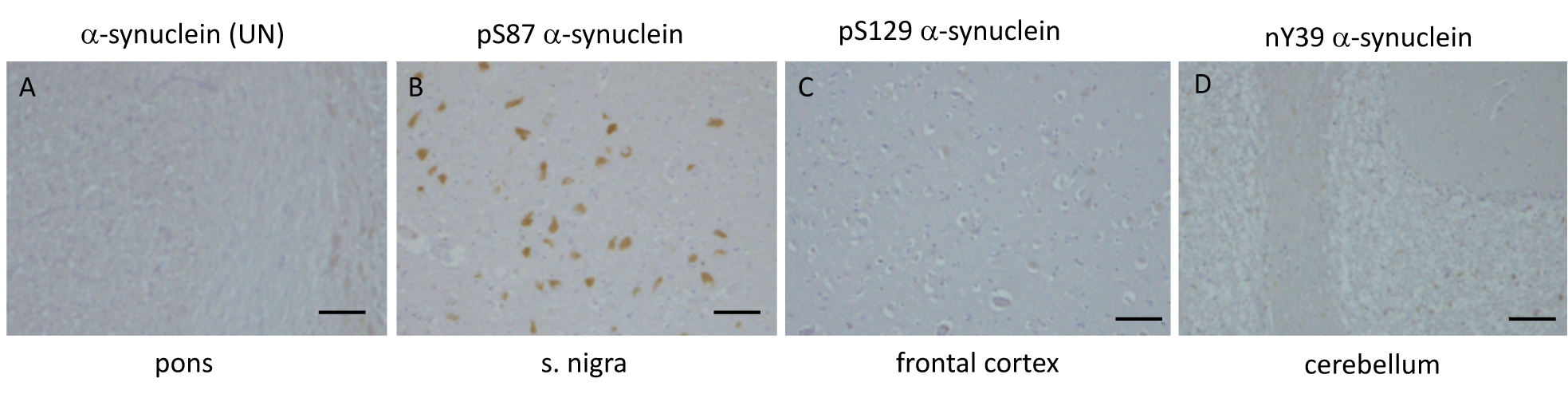

**Figure S1.** Representative immunohistochemistry illustrations of neurologically normal control cases showing no abnormal α-synuclein deposits when treated with various α-synuclein antibodies. Scale bar 50μm in all.

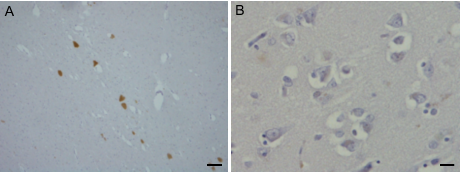

**Figure S2.** Immunohistochemistry of brain sections from IPD cases showing no appreciable staining within PD nigra (A) or frontal cortex (B) when primary antibodies were omitted but treated with anti-rabbit and anti-mouse secondaries respectively. Scale bar 60μm in A, 15μm in B.
